# Reconstruction of segmental duplication rates and associated genomic features by network analysis

**DOI:** 10.1101/2023.03.18.533287

**Authors:** Eldar T Abdullaev, Dinesh A Haridoss, Peter F Arndt

**Affiliations:** Department of Computational Molecular Biology, Max Planck Institute for Molecular Genetics, Berlin, Germany; Department of Electrical Engineering and Computer Science, Indian Institute of Science Education and Research Bhopal, Bhopal, India

**Keywords:** Segmental duplications, Complex networks, High-copy repeats

## Abstract

Segmental duplications (SDs) are long genomic duplications fixed in a genome. SDs play an important evolutionary role: entire genes together with regulatory sequences can be duplicated. Ancestral segmental duplications gave rise to genes involved in human brain development, as well as provided sites for further genomic rearrangements. Some duplicated loci were extensively studied, however, universal principles or biological factors of SDs propagation are not fully described yet. Segmental duplications can be arranged into a network where edges correspond to real duplication events, while nodes to affected genomic sites. This gave us an opportunity to estimate how many duplications happened in each locus. We studied genomic features associated with increased duplication rates with especial interest in high-copy repeats distribution relative to duplicated regions breakpoints. Our comprehensive study of genomic features associated with duplications and those associated with increased duplication rates allowed us to identify several biological factors affecting a segmental duplication process. We found genomic features associated with increased duplication rates, three signatures of duplication process and associations of SDs with different classes of high-copy repeats.

## 1 Background

Segmental duplications (SDs) or low-copy number repeats are long duplications of genomic sequence that are fixed in a population. Conventionally, only those duplications longer than 1 kbp and with the level of sequence identity > 90% are annotated as segmental duplications. Assuming a clock-like mutation rate in the human genome, the SDs defined this way correspond to duplication events that happened after the divergence of the New and Old World monkeys Bailey and Eichler [2006] She et al. [2006] about 40 Myr ago. Based on the relative position of the 2 copies of SDs they are classified as intrachromosomal (when both copies are located on the same chromosome) or interchromosomal (when those are on different).

SDs are distributed very non-uniformly in the human genome. Some loci are extensively duplicated while other are depleted of segmental duplications. For example, pericentromeric and subtelomeric parts of chromosomes are enriched with SDs, especially with interchromosomal ones Linardopoulou et al. [2005], She et al. [2004], Guy et al. [2003]. General principles of segmental duplications propagation in the human genome were suggested when SDs were studied as a graph Abdullaev et al. [2021]. A specific SD network where nodes represent genomic loci involved in duplications, while edges correspond to alignments between those loci was constructed and it’s growth was modelled. As a result, a model where duplication rates preferentially grow with node degrees of corresponding nodes (the preferential copying model or PCM) is the one according to which segmental duplications are accumulated in the human genome and probably genomes of other vertebrate species Abdullaev et al. [2021]. Other than this rather universal dynamical principle of SDs evolution there are many genomic characteristics that seem to affect duplication rates.

A slight positive correlation of segmental duplications distribution with the gene density was reported by Zhang et al. [2005] along with negative correlation with recombination rates. Large duplications tend to be enriched in heterochromatic parts of the genome or, according to other observations, in hetero- to euchromatin transition regions Grunau et al. [2006], Kirsch et al. [2008]. Duplicated regions are, on average, of higher G/C content in comparison with the rest of the genome Bailey et al. [2003], Zhang et al. [2005]. In agreement with that, copy number varinats (CNVs) breakpoints seem to be enriched with G/C-rich sequences predicted to form G-quadruplexes Bose et al. [2014]. The listed factors are observed irregularly and not necessarily reproduced in other experimental settings. Also these associations can be specific to some chromosomes, but absent in others Zhang et al. [2005].

High-copy number repeats can cause segmental duplications through homology-mediated mechanisms. Thus some repeat classes are enriched at breakpoints of SDs since they provide necessary stretches of homologous sequences. Some cases of duplication causing repeats were reported even before SD annotation in the human genome Eichler et al. [1996, 1999], Guy et al. [2000]. The systematic study by Bailey et al. [2003] identified those repeat classes that are significantly overrepresented at SDs breakpoints when compared with the genome average. Only those accurately annotated SDs not overlapping other duplications were analyzed. Two repeat classes are significantly overrepresented at breakpoints: *Alu* repeats and satellites (specifically, *HSATII, GSAT,* and *TAR1*), while many repeat classes are even underrepresented in flanking regions. Specifically, among Alu subfamilies, younger ones (*AluY* and *AluS*) account for the enrichment, while the oldest primate subfamily (*AluJ*) does not Bailey et al. [2003]. The burst of *Alu* retroposition around 35-40 million years made an ancestral human genome more susceptible to segmental duplications Bailey and Eichler [2006], Bailey et al. [2003]. Later resulting SDs themselves became a source of homology for further duplication events.

Replication timing is another factor associated with duplications. For example, it was observed that recurrent CNVs are enriched in early replicating genomic sites, while non-recurrent ones are in late replicating regions Koren et al. [2012], Chen et al. [2015]. Other than that, some groups observed enrichment of CNVs in early replicating regions Lu et al. [2014], while other in late ones De and Michor [2011]. Finally, it was suggested that division into early and late replicating regions is not sufficient to explain CNVs distribution. The hotspots of CNVs are significantly associated with sites of reduced DNA polymerase velocity (those where the difference in replication timing is the largest), in other words, in transition regions located between early and late replicating sites Chen et al. [2015].

In this study we are focused on finding genomic features affecting duplication rates. To solve this task we exploit the network representation of human SDs which we described above. Firstly, based on characteristics of the SD network we approximately evaluate the number of duplication events that each duplicated region (or node of the SD network) underwent in the past evolution. Then we analyze associations of genomic features with duplication rates for those regions. Finally, we looked at characteristic patterns or signatures of SDs.

## 2 Results

### 2.1 The SD network construction

In our analysis we used annotated SDs of the human genome Bailey et al. [2001, 2002]. The annotation includes a list of pairwise alignments between regions of the GRCh38 reference genome which are longer than 1 kbp with at least 90% of sequence identity between copies. Not all such alignments correspond to actual duplication events, some of them appear as a result of overlap between an already duplicated region and a new SD (Fig. 1). As a result a new copy is homologous to both an ancestral region and its another copy. However, only the first of two alignments correspond to an actual duplication event. We call this type of alignments “primary” as opposed to “secondary” alignments which appear because of overlaps between duplications and not correspond to a duplication event between a pair of sequences.

**Figure 1:**
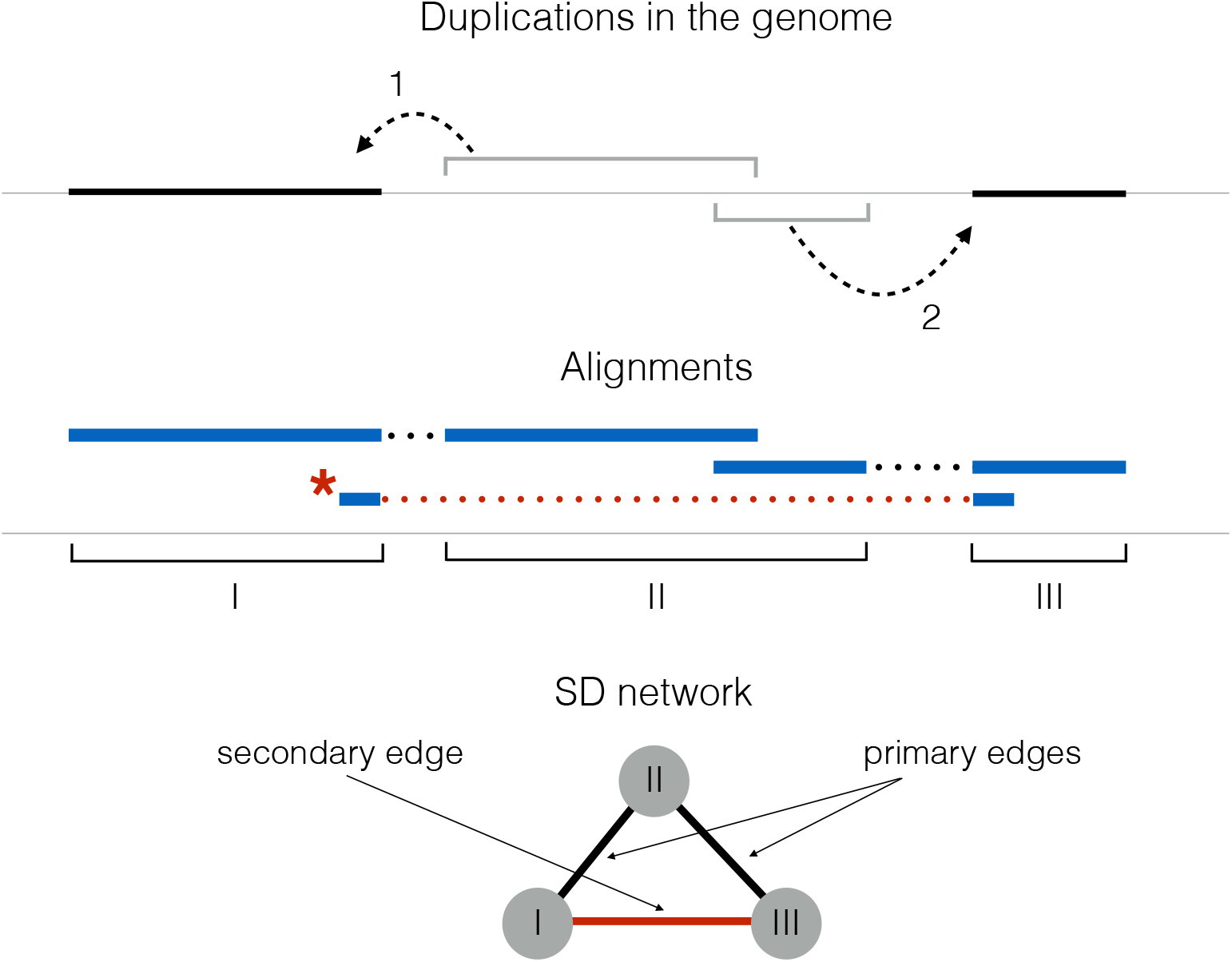
A simple example of two duplication events in a genome and an SD network that corresponds to them. Duplication events are marked with arabic numerals while resulting groups of overlapping alignments (duplicated regions) with roman ones. A duplicated region can appear as a result of one or several duplication events and is not overlapping any other duplicated regions. Our two duplications result in two primary and one secondary (marked with the red asterisk) alignments. The last one does not correspond to a duplication event and appears as a result of overlapping duplications. The SD network is constructed in the following way: nodes represent duplicated regions (marked with respective roman numerals), edges are added if an alignment between two duplicated regions exists.

From all human autosomal SDs we generated the network of segmental duplications or the SD network as described in Abdullaev et al. [2021]. Each node of the network represents a duplicated region, which we define as a genomic interval that covers a maximal set of overlapping alignments. In other words, a duplicated region equals to a union of all overlapping alignment intervals. Undirected edges are added if alignment between two corresponding duplicated regions (nodes) exists (Fig. 1). The resulting SD network includes 6656 nodes and 16,042 edges which are organized in 1999 connected components. Among those components one stands out in the size. The giant component contains 1325 nodes (19.9% of all nodes) and 9678 edges (60.3% of all edges).

As described above, edges of the SD network represent alignments of homologous sequences that share common origin, however, the information on the temporary order and direction of duplications (i.e. which genomic region among two copies is ancestral) is missing. Moreover, edges of the SD network represent either real duplication events or “secondary” alignments that appear because of overlaps between independent duplications. Reconstruction of real duplication events from the whole network of duplicated regions would allow us to further look into biological factors responsible for segmental duplications.

### 2.2 Duplication event reconstruction. Graphs without cycles

Before going into our method of duplication events reconstruction, one has to understand the nature of cycles in the SD network. Given the fact that the SD network evolved according to the preferential copying model we can do some speculations about those cycles. Cycles in PCM networks appear as a result of duplication when a new “daughter” node inherits an edge to a neighbor of the “mother” node from it (Sup. Fig. 1a). Only cycles of size 3 can appear as a result of such secondary edges acquisition in PCM. Indeed in a cycle of size *K* where *K* > 3 we expect at least one event of secondary edge acquisition (as the only way to get a cyclic structure). On the other hand, it is impossible to acquire a neighbor from the “mother” node that does not have one (Sup.Fig. 1b), thus only cycles of size 3 appear as a result of node duplications in the PCM. We analyzed cycles and shortest self-paths (shortest path from a node to itself where exists) of the SD network. In agreement with this we observe that our SD network is depleted with cycles of size > 3 and shortest self-paths longer than three edges in comparison with other networks of well-known topology (Table 1).

**Table 1:**
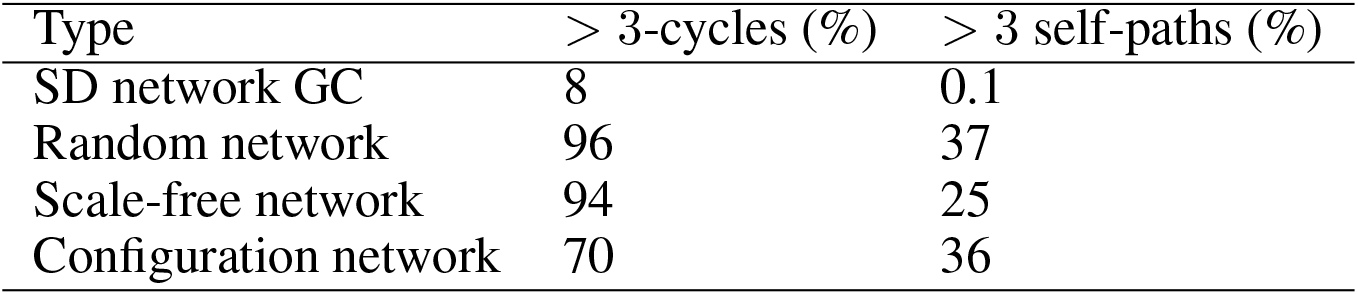
The characteristics of several networks that include: random graph, scale-free network, the giant component of the SD network and corresponding configuration network (random graph with node degrees of original network preserved). All considered networks are of the same number of nodes. The columns are: the fractions of > 3 nodes cycles and > 3 edges self-paths. The SD network is strongly depleted with large cycles in comparison with the other considered networks. This difference is even more prominent when considering self-paths (< 0.1% of all self-paths). This depletion supports the earlier prediction that the SD network evolved according to one of copying models. Larger cycles that we still observe in the SD network, likely, originate from superposition of several size 3 cycles (Sup. Fig. 1c).

| Type | $> 3$-cycles (%) | $> 3$ self-paths (%) |
| --- | --- | --- |
| SD network GC | 8 | 0.1 |
| Random network | 96 | 37 |
| Scale-free network | 94 | 25 |
| Configuration network | 70 | 36 |

Let’s consider a case of duplication events reconstruction from an example duplication network without any cycles. If we disallow any inheritance of neighbors from the “mother” node (e.g. by simulating PCM with *f* = 0) we will observe a network without any cycles where each edge represents a real duplication event. Directionality of duplication events can be reconstructed in a unique way if we assume that we know which node was an ancestral one for each connected component (Fig. 2). By comparing two figures (Fig. 2a and Fig. 2b) we can see that directionality of edges changes when taking another ancestral node while the number of duplications that each node underwent is almost invariant to this choice. Thus for our task it is sufficient to find edges in the SD network that correspond to real duplications while assigning directionality is not needed. In other words, resolving cycles in the SD network by excluding all secondary edges would be enough to say how many times each node duplicated in the course of the network growth.

**Figure 2:**
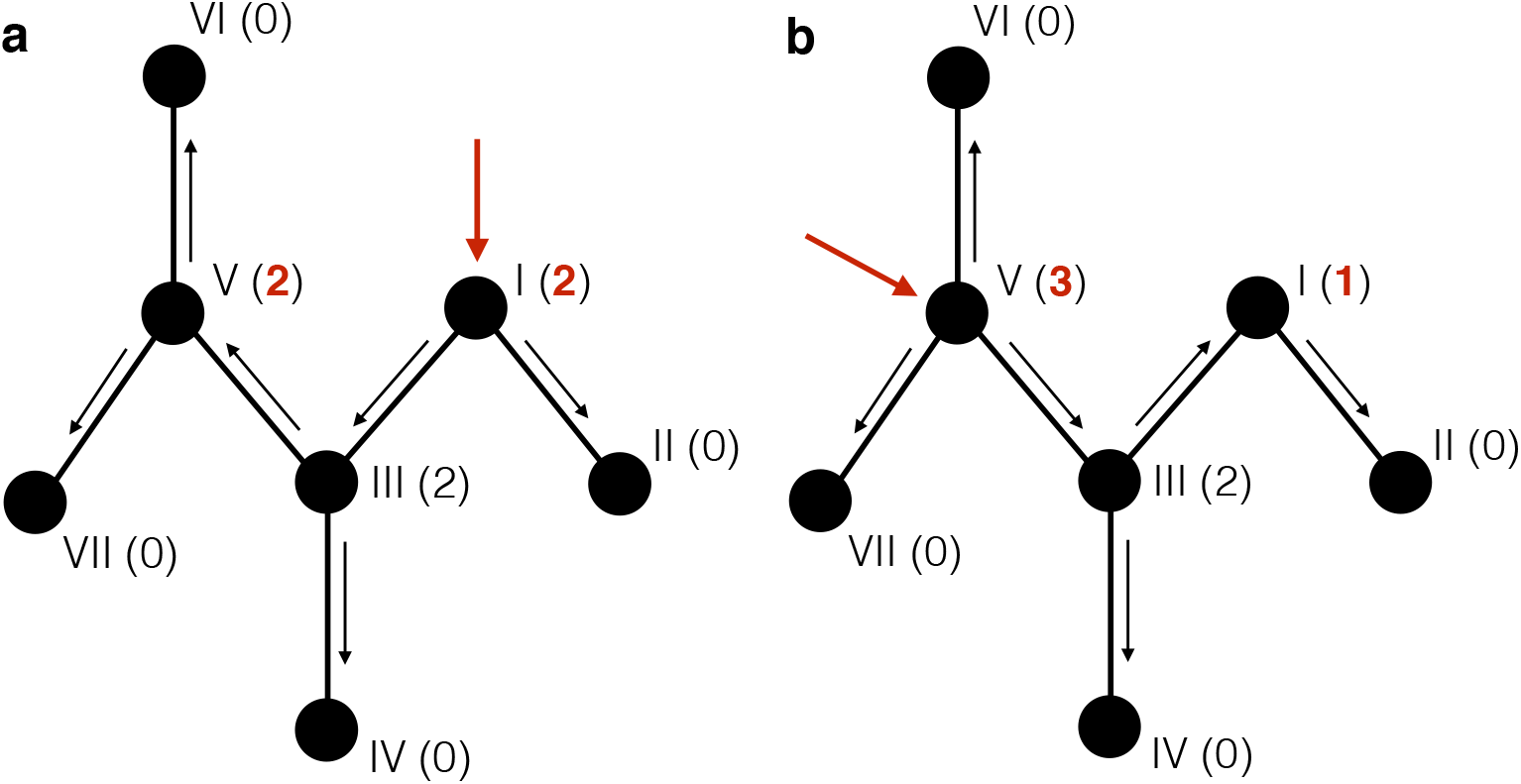
**a**. The reconstruction of duplication events is a straightforward task when cycles are absent. Assuming we know *a priori* which node was the first in a simulation (pointed by the red arrows), there is a single reconstruction of events possible. One can reconstruct all “mother” to “daughter” nodes relationship, however, information on temporary order can not be inferred this way. The numbers in parentheses denote the number of duplications each node made in a course of a network growth. Assigning another starting node (**b**) results in distinct pattern of duplications, however, overall number of duplications each node made stays almost intact. The numbers differ only for suggested starting nodes. In practice, this means that to get an information on how many times each node of the SD network duplicated - one has to distinguish edges representing real duplication events (primary edges) from secondary ones. There is no need in assigning directionality or suggesting the first node for our task.

### 2.3 Duplication event reconstruction. Graphs with cycles

The only way how cycles of size three can appear in a PCM generated network is when a daughter node inherits an edge from a mother node. This can happen in several scenarios (Fig. 3a) so reconstruction of duplication events from a network with cycles is not a trivial task anymore. It can be formulated as a search for a subgraph tree (a spanning tree) that goes through all nodes of the SD network and covers only those edges that correspond to real duplication events (Fig. 3b). The spanning tree by definition does not have any cycles, thus for the SD network with *N_STree_* = 6656 nodes, *E_SD_* = 16, 042 edges and *C_SD_* = 1999 components we expect a spanning tree with *N_STree_* = *N_SD_* = 6656 nodes, *E_STree_* = *N_SD_* – *C_SD_* = 4657 edges and *C_STree_* = *C_SD_* = 1999 components. There are several existing algorithms that can find the minimum spanning tree (MST) for a graph that goes through edges of minimal overall weight. To apply the algorithms one has to assign weights to the edges of the SD network, such that lower weights correspond to likely primary edge. We use an heuristic approach to assign such weights. This approach is based on a set of assumptions. First of all, we study each node of the SD network independently. Secondly, according to the PCM each node of a connected component except for the ancestral pair of nodes appears as a result of duplication of its mother node. Moreover, a mother node has to be among the neighbors of a node of interest, because according to the PCM the only guaranteed edge of a daughter node is the one with a mother node. Finally, it is expected that a daughter node shares the highest fraction of its neighbors with a mother node. With these three assumptions we can assign edge weights as further detailed in the corresponding section of the Supplementary materials.

**Figure 3:**
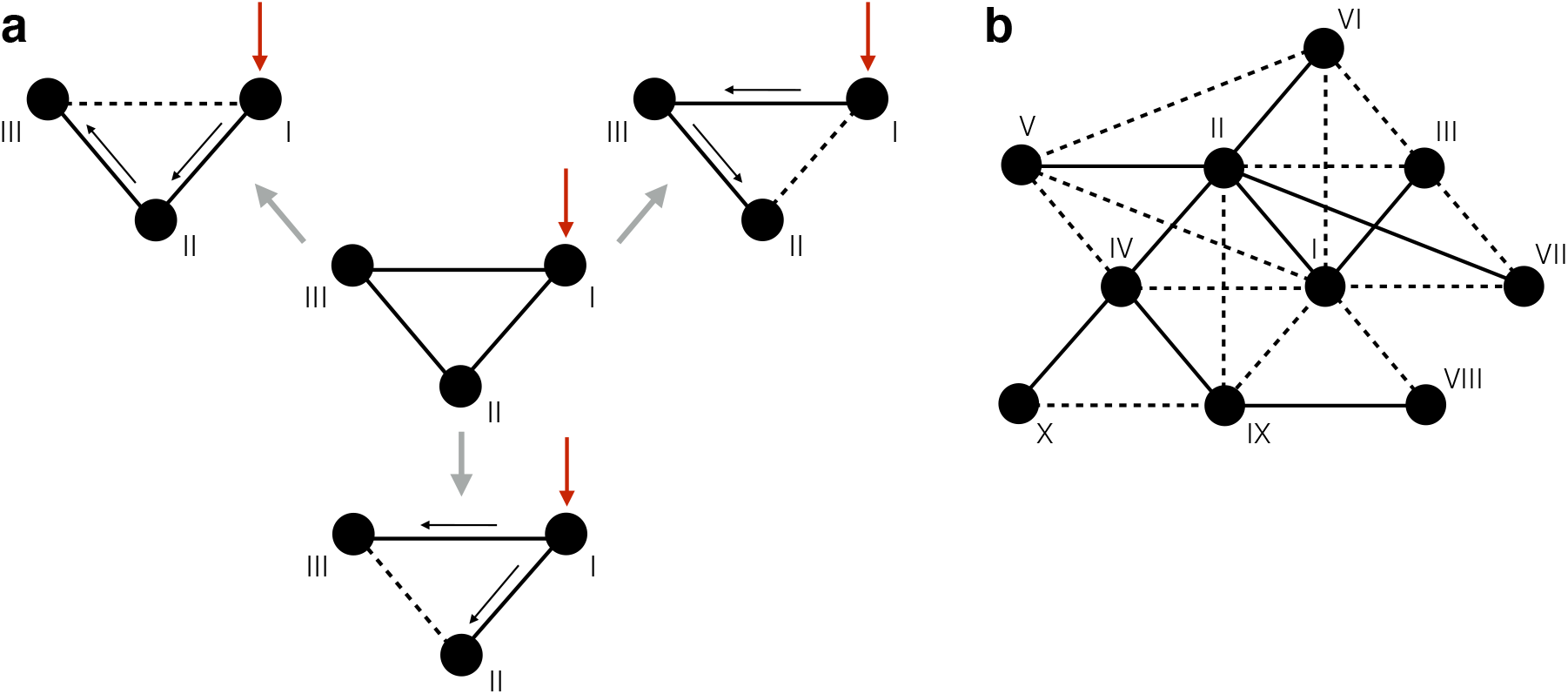
Reconstruction of duplication events in graphs with cycles. The black arrows represent duplication events (pointing from “mother” to “daughter” nodes), red arrows represent suggested starting nodes, the dotted lines represent secondary edges. One can see that the cyclic graph in the center of the scheme (**a**) can be reconstructed in three different ways even when the first node stays the same (because each of three edges in the cycle can be secondary). The number of possible configurations grows with the number of secondary edges (or cycles) in the network of interest. The scheme (**b**) illustrates one possible reconstruction of duplication events in a more complex network. The subgraph of reconstructed duplications has to be of a tree structure (connected and acyclic).

In practice, our approach was implemented as the following weighting scheme. For each edge *e* of the SD network connecting nodes *a* and *b* we assigned the weight 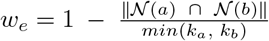, where 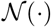 is a set of neighbor nodes of a node of interest, || · || denotes the number of elements in a set and *k* is a node degree. Values *w_e_* lie in the interval [0 < *w_e_* ≤ 1] where *w_e_* = 1 when no neighbors are shared, while *w_i_* > 0 because there is always at least one neighbor not included in the overlapping set 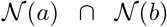. In other words, 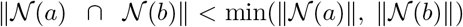 since *a* is not in *N*(*a*) and *b* is not in *N*(*b*) by definition. As a result, edges connecting nodes with many shared neighbors are of lower weight and thus more likely being included in the minimum spanning tree. These edges are of our interest, because likely correspond to real duplication events.

### 2.4 Accuracy evaluation

We tested the accuracy of our approach on PCM simulated networks. Two metrics were used to measure the accuracy of our predictions: the fraction of correct (primary) edges in the MST and a variance explained in duplications number that each node underwent during the simulation (Table 2). We compared our method, listed as “Predicted MST”, with several alternative strategies of predicting number of duplication events per node (listed in the Table 2). These include multiple node centrality measures. We also considered two trivial models: simple use of PCM network node degrees and a random spanning tree. These models define baseline accuracy levels. One can see that our heuristic approach performs the best among other methods and explains 77% of variance in number of duplications while trivial use of node degrees reaches only 58%. It is illustrative to the fact that the reconstruction step we made leads to a better approximation of the number of duplications than a trivial use of node degrees.

**Table 2:** Comparison of several methods for predicting the number of duplications that each node underwent during PCM simulations. To evaluate an accuracy we measured the fraction of correctly predicted primary edges (where possible) and a variance explained for the vector of real duplications number (*R*^2^). We measured an accuracy of our MST based algorithm described in the main text (“Predicted MST” row in the table). We used multiple centrality measures for nodes in resulting PCM synthetic networks as listed; one can find more information about these measures Seth Bromberger [2017], Brin and Page [1998]. We considered the vector of node degrees and a random spanning tree covering nodes of a synthetic network as a trivial models to estimate a baseline quality of predictions. One can see that our algorithm performs the best in both criteria among all alternatives with, per average, 64% of primary edges correctly predicted and 77% of variance in number of duplications explained.

| Method | Edges match (%) | Variance explained ($R^2$) |
| --- | --- | --- |
| Predicted MST | 64 | 0.77 |
| Node degrees vector | *NA* | 0.58 |
| Random spanning tree | 20 | 0.44 |
| Betweenness (edges) | 17 | 0.64 |
| Betweenness (nodes) | *NA* | 0.66 |
| Closeness centr. | *NA* | 0.17 |
| Eigenvector centr. | *NA* | 0.48 |
| PageRank centr. | *NA* | 0.67 |
| Radiality centr. | *NA* | 0.02 |

Then our approach was applied on the real SD network. From the resulting MST that covers putative duplication events in the SD network we calculated the number of duplications that happened in each node. Since edges in the resulting MST represent duplication events we can say that almost every node *i* (except for the first one) in the network was duplicated *k_i_* – 1 times where *k_i_* is a node degree of *i^th^* node in the MST.

To validate the accuracy of our prediction we further analysed characteristics of edges predicted as primary duplications. For example, matching coordinates of a pair of SDs is a characteristic feature of a secondary alignment. These “suspicious” alignments often appear as a result of overlap between duplication events (see Sup. Fig. 2a and Supplementary Materials). We observed that “suspicious” edges with matching breakpoints were depleted in our predicted MST when comparing with random samples of edges from the rest of the network (empirical p-value < 0.001). Secondly, we expect that a correct MST covering real duplication events includes more edges of higher sequence identity than a random set of edges from the SD network. We found that our predicted MST for the SD network is enriched with highly identical edges (alignments). There are more alignments of sequence identity higher than 0.99 in the predicted MST in comparison with random samples of edges from the rest of the SD network (empirical p-value = 0.001). We explained this effect in more detail in the Supplementary Materials and Sup. Fig. 2b.

### 2.5 Associations with genomic features

#### 2.5.1 Genomic features associated with duplicated regions

For each duplicated region (node) of the SD network we collected multiple genomic features associated with that region. These genomic features include: replication timing, recombination rates, openness of chromatin, genome assembly gaps, CTCF binding sites, coordinates of a duplicated region, G/C nucleotides content, number of gene overlaps, repeat overlaps and CpG island overlaps (see Table 3 for details). Some features were measured inside the duplicated regions (between the breakpoints), while some in flanking regions of length 50 bps (“Position” column at Table 3). We used those features to find associations with the number of duplications that we predicted during the reconstruction step. In other words, we formulated a machine-learning task of predicting the number of duplications (response variable) given the matrix of genomic features associated with corresponding duplicated regions (predictor variables).

**Table 3:** The columns “Type” and “Position” specify how and where a feature was measured (either inside of a duplicated region or in two flanking regions of 50 bps). For cell-line specific features we used either a data from the GM12878 lymphoblastoid cell line (picked arbitrary) or the master track for DNAse hypersensetivity. The master track represents an integrated DNAse hypersensetivity data for 125 separate cell lines (one can read more on that at http://genome.ucsc.edu/cgi-bin/hgTrackUi?db=hg19&g=wgEncodeAwgDnaseMasterSites and Thurman et al. [2012]).

| Genomic feature | Type | Position | Cell line |
| --- | --- | --- | --- |
| Coordinates and length | value | — | — |
| Telo-/centromere dist. | value | — | — |
| Intrachrom. edges (%) | fraction | — | — |
| Replication timing | mean | inner + flanks | GM12878 |
| Replication pausing | span | inner + flanks | GM12878 |
| Recombination rate | mean | inner + flanks | — |
| DNase hypersens. | mean | inner + flanks | master (125 cell types) |
| Assembly gaps | count | flanks | — |
| CTCF sites | count | flanks | — |
| G/C content (%) | fraction | inner + flanks | — |
| Genes | count | inner | — |
| CpG islands | count | inner + flanks | — |
| DNA transposons | count | flanks | — |
| LTR retrotransposons | count | flanks | — |
| LINEs | count | flanks | — |
| SINEs | count | flanks | — |
| Satellite repeats | count | flanks | — |

Firstly, we studied if various characteristics of duplicated regions differ from those observed at random genomic sites. We compared duplicated regions against the genome background distribution without, as for now, taking into account the number of duplications that they underwent. Multiple characteristics of duplicated regions were either significantly larger or smaller than expected from the null hypothesis of random distribution (Fig. 4). It is not surprising given the fact that duplications are distributed non-uniformly in the human genome. We observed that assembly gaps are enriched at flanking regions (because large complex duplications are often hard to properly map), duplicated regions are located in late replicating regions or/and in those where DNA polymerase slows down, recombination rates are lower in duplicated regions, while the G/C content is higher.

**Figure 4:**
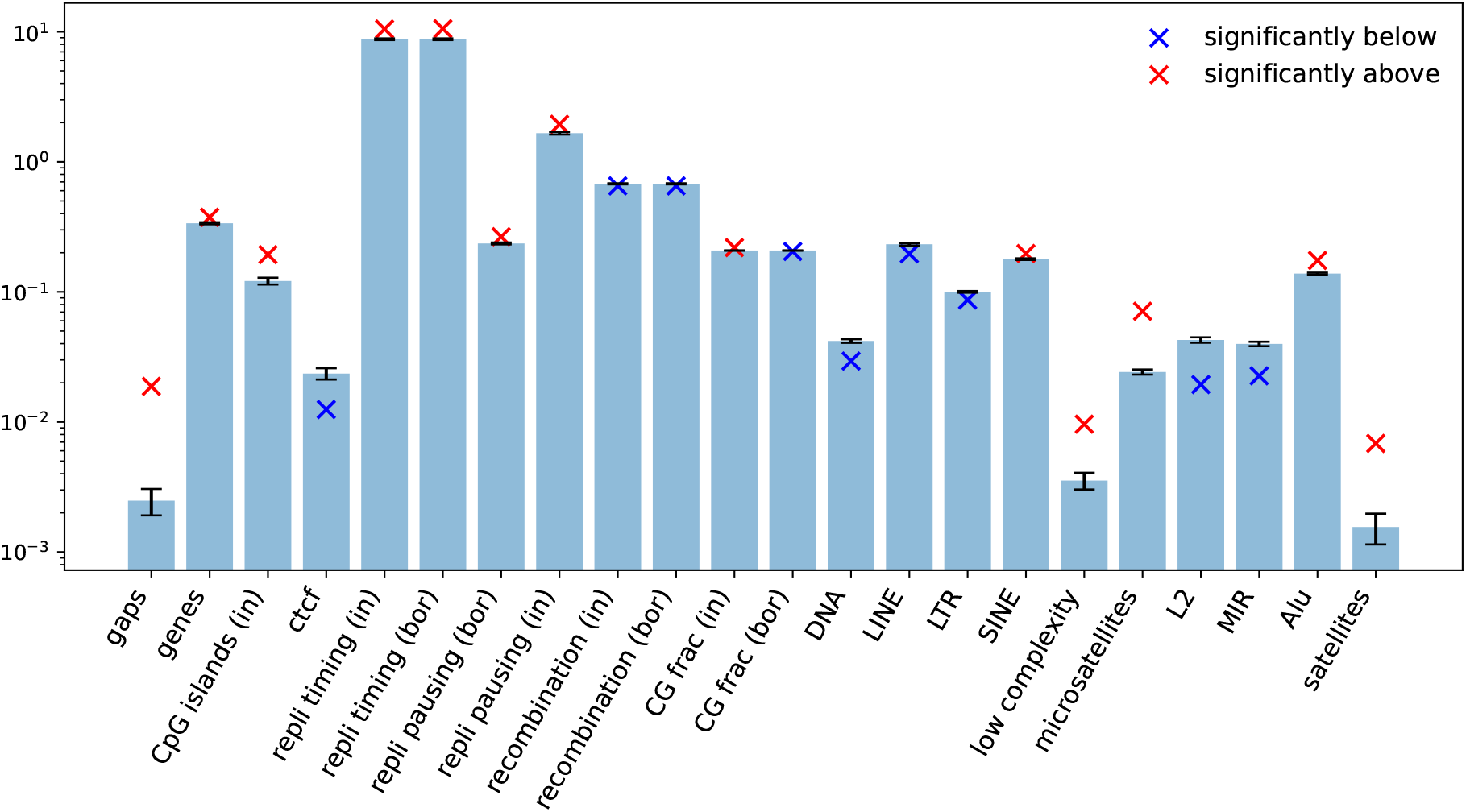
Characteristics of duplicated regions in comparison with the genome average. The barchart represents distributions of various genomic features (listed on the *X* axis) measured in randomly shuffled genomic intervals. The crosses are values that we observe for original duplicated regions (red ones are above and blue ones are below the background mean). Only those features where the difference is significant are listed out of the whole list of features.

**Figure 5:**
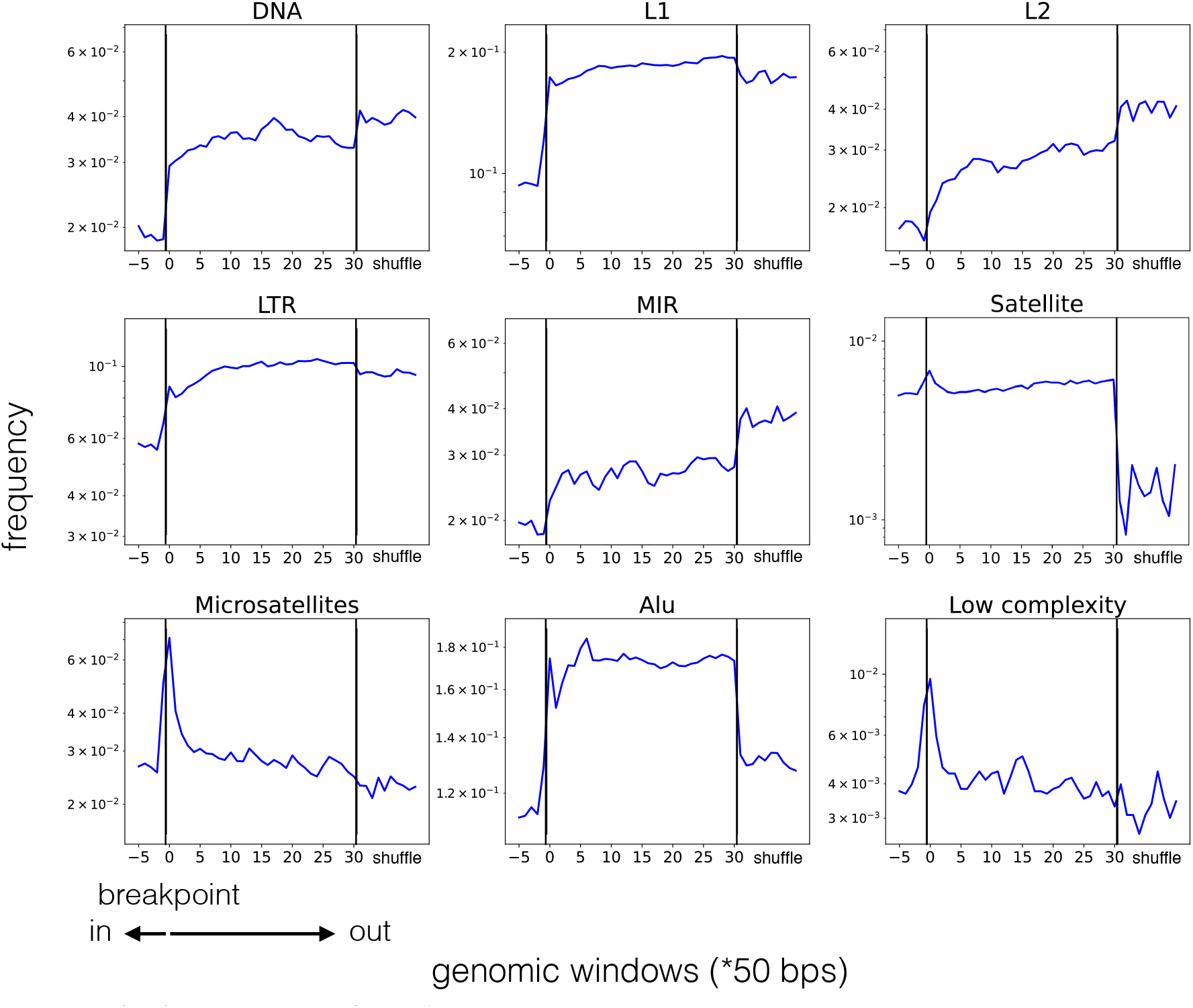
The distribution of different repeat families relative to the breakpoints of duplicated regions. Technically, we took two genomic windows of 50 bps upstream and downstream of all duplicated regions, calculated the number of repeats falling into those windows and divided it by the overall number of those windows (the frequency values on the *Y* axes). We did 5 measurement steps inside of duplicated regions, 30 measurements outside and 10 times in random genomic positions. The *X* axes represent genomic positions relative to breakpoints (marked with vertical line); last 10 points separated by another vertical line correspond to counts in randomly placed genomic windows. The first group of repeats (from DNA transposons to *MIR* repeats) includes those depleted inside of duplicated regions with no enrichment observed around the breakpoints. The second group (from satellites to low complexity repeats) includes repeats that are enriched at the very breakpoints of duplicated regions or in nearby windows outside (satellites, microsatellites, *Alu* and low complexity repeats).

We studied in detail how high-copy repeats are distributed relative to duplicated regions breakpoints (inside, outside or at borders of duplicated regions). To do this we moved sliding non-overlapping windows of length 50 bps inside and outside of breakpoints of duplicated regions and counted the number of overlapping repeats (see Methods for details). Also for each repeat class we did 10 more measurements in random genomic positions. As a result, the distributions were different for different repeat classes. These could be divided into two groups. Repeats in the first group include: DNA transposons, *LTRs, L1* and *L2* repeats from the LINE family and *MIR* repeats from SINE. These are depleted inside of a duplicated region and their numbers grow when we move further outside from a duplicated region borders (Fig. 7). Repeats in the second group are enriched in a close proximity outside of duplicated regions, especially at the very breakpoints. This group includes: satellites, microsatellites, *Alu* and low complexity repeats (illustrated at Fig. 7 subplots, starting from the satellites on). The low complexity repeats are composed of polypurine or polypyrimidine repeated stretches, or regions of high A/T or G/C content. The second group of repeats is likely responsible for genomic instability that leads to duplication events. It is unlikely that observed biases are attributed to a non-uniform SDs distribution in the genome (concentration of SDs in subtelomeric or subcentromeric regions, proximity to assembly gaps etc.). Firstly, because repeats from the second group are depleted inside of duplicated regions and at a relatively small distance from them and, secondly, because exclusion of duplicated regions flanked by the assembly gaps did not change the observed distributions. The difference in repeats frequency between proximal to breakpoints and background genomic windows is significant for all repeat families, except for *L1* (Fig. 4).

#### 2.5.2 Prediction of genomic features associated with increased duplication rates

We trained a random forest model (RF) to find genomic features associated with the number of duplications a region underwent. We estimated the feature importance that is assigned to predictor variables by the RF algorithm with additional rounds of permutations (see Methods or Altmann et al. [2010]). As a result, the following genomic features were important in the number of duplications prediction: length of a duplicated region (emp. p-value < 0.001), fraction of intrachromosomal edges (emp. p-value < 0.001), number of overlapping genes (emp. p-value = 0.023) and CpG islands (emp. p-value = 0.0025), replication pausing (emp. p-value = 0.025) (the last two measured inside of a duplicated region). Significant importance of those features cannot be attributed to the fact that some of those features are correlated with each other. To clarify the type of dependence between the features and response variable we calculated partial Spearman’s correlation coefficients for each predictor variable (genomic feature) with the response variable (number of duplications underwent) controlling for all other predictors as possible confounding variables (Fig. 6). Additionally, we measured the regular Spearman’s rank correlation coefficient and estimated its significance by permutations. One can interpret these coefficients in the following way: a regular correlation coefficient represents the dependence we would observe without considering other correlated features. However it could be a result of other confounding correlations that are taken into account when the partial correlation is calculated. It shows the effect of one variable on another independent of other variables.

**Figure 6:**
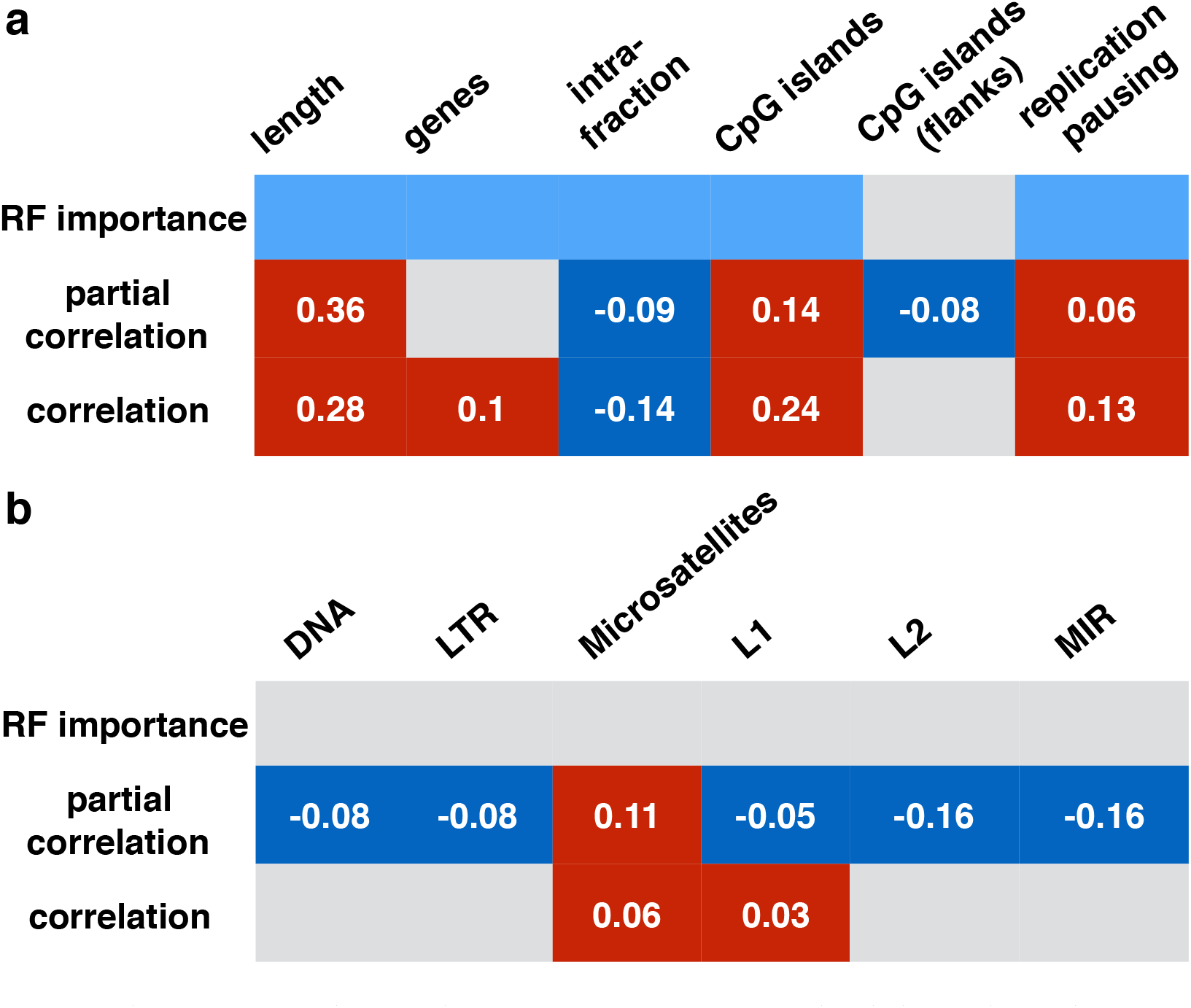
Characteristics of duplicated regions significantly associated with the number of duplications. Only significant associations predicted by the random forest, partial or regular Spearman’s correlations are listed. **a**. These include: length of a duplicated region, fraction of intrachromosomal edges from all edges of a node, number of CpG islands overlapping a duplicated region or its flanks, replication pausing and number of overlapping genes. The cells coloured in light blue represent those features with high permutation importance in the random forest algorithm. Dark blue and red cells correspond to features with significant negative or positive correlation coefficients (written inside of the cells) while unfilled cells represent non-significant ones. **b**. High-copy repeats that show non-zero partial correlation coefficients with the number of underwent duplications. Most of those associations are not supported by the random forest or significant non-zero Spearman’s correlation coefficient.

**Figure 7:**
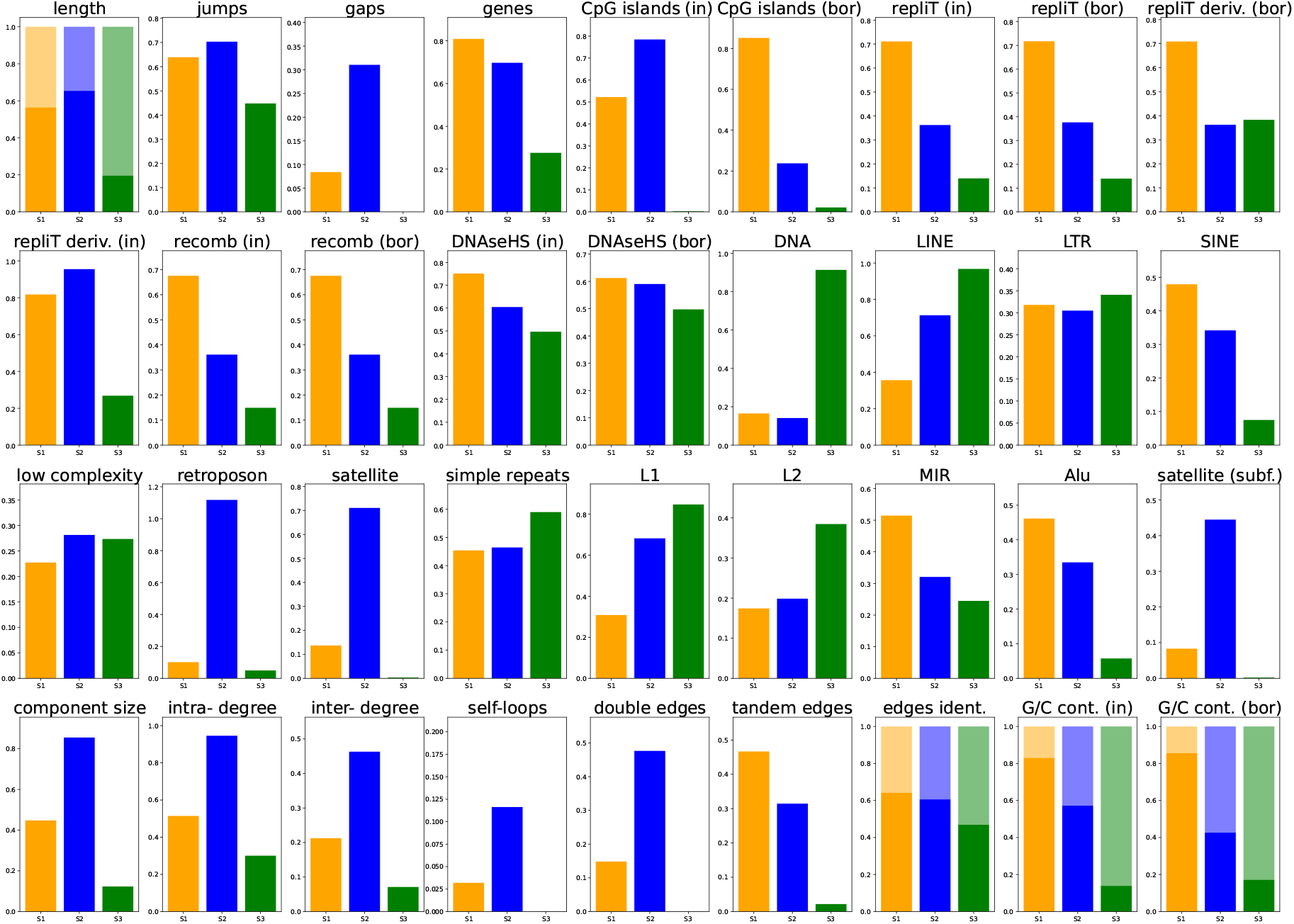
Characteristics of signatures s1, s2 and s3 colored in orange, blue and green, respectively. Four features (length, sequence identity, G/C content inside and at flanks) are binarized to below (lighter) and above (darker color) their median value.

The partial and regular Spearman’s correlation coefficients are significant and positive for the length of a duplicated region, number of CpG islands (inside) and replication pausing (inside), while negative for intrachromosomal edges fraction. Other correlation coefficients are not significantly different from zero. As we mentioned earlier, the loci with reduced DNA polymerase velocity could be associated with genomic duplications Chen et al. [2015]. Our results support this hypothesis: the association of replication pausing and duplication frequency is reproduced by several our approaches (Fig. 6a). On the other hand, we did not find evidence of other associations with late or early replicating regions. Association with the number of gene overlaps also makes biological sense since segmental duplications were responsible for gene families propagation and evolution in the human lineage. To our knowledge, only increased G/C content was observed in segmental duplications, however, enrichment of CpG islands in highly duplicated regions is not described yet. Several high-copy repeat families had significant non-zero partial correlation coefficients (Fig. 6b). However, most of these associations were not supported by significant Spearman’s correlation coefficients or the RF importance estimates.

### 2.6 Analysis of segmental duplication signatures

We considered a scenario where several mechanisms of SDs accumulation act in the genome. Each one results in SDs of different genomic characteristics. We applied non-negative matrix factorization or NMF to find signatures of duplication processes (see Methods). NMF is a method of approximate matrix factorization into product of two matrices with all non-negative elements. These matrices are usually interpreted as signatures matrix (one with characteristic patterns of some process) and weights matrix (with impacts of each signature at each sample) Paatero and Tapper [1994]. This method allows to detect characteristic patterns of SDs accumulation if present, without any *a priori* knowledge about duplication mechanisms. We applied NMF to the matrix of duplicated regions characteristics in 10 Mb genomic windows. For each genomic window we took duplicated regions that overlap it, extracted their genomic features (see Table 3) and summed them up. Some features were binned relative to median before summing them up. These include: length, G/C content and sequence identity. We applied NMF on resulting matrix. To predict the optimal number of signatures we used several approaches listed in the Methods section (Sup. Fig. 4). As a result, NMF with s = 3 signatures performed the best. These are three patterns of SDs accumulation which we observe in the human genome. We will further denote them as signatures s1-s3.

One can see that the signature s2 corresponds to long duplications with high node degree which tend to lie near assembly gaps (hard to assemble multi-copy regions) and dominate the giant component of the SD network (Fig. 7). Two more duplication patterns were predicted for shorter duplicated regions. The signature s1 represents SDs located in late replicating regions of higher recombination rates and G/C content in comparison with those harboring s3 duplications. Moreover, s1 duplicated regions overlap genes more often than s2 and s3 ones despite the fact that s2 duplicated regions are longer. This means that s1 signature events are responsible for gene duplications. All 3 signatures differ in major repeats representatives at their flanks. The signature s1 is dominated with SINE (both *MIR* and Alu) repeats, s2 is dominated with non-autonomous retrotransposons and satellite repeats, while s3 with DNA transposons and LINE (both *L1* and *L2*) repeats. We assigned one of our three signatures to each individual duplicated region as described in Methods section. As a result we got 2510, 704 and 3442 duplicated regions assigned to s1, s2 and s3, respectively.

We observed that duplicated regions belonging to the various signatures are distributed non-randomly relative to telomeres and centromeres (Fig. 8). Both s2 and s3 duplications are enriched in subtelomeric and pericentromeric regions while their numbers drop as we move away from them. s1 events, on the other hand, are distributed more uniformly with no enrichment around centromeres or telomeres. Overall, these observation mean that s1 duplications likely originate from SINE-based non-allelic homologous recombination which happens in interstitial parts of chromosomes. It agrees with the fact that this type of events are usually *Alu* dependent. Also one characteristic scenario of NAHR involves misalignment of *Alu* repeats with consequent tandem duplication of genomic locus between those repeats. We suggest that the highest rate of tandem events in s1 duplications can be explained by this scenario.

**Figure 8:**
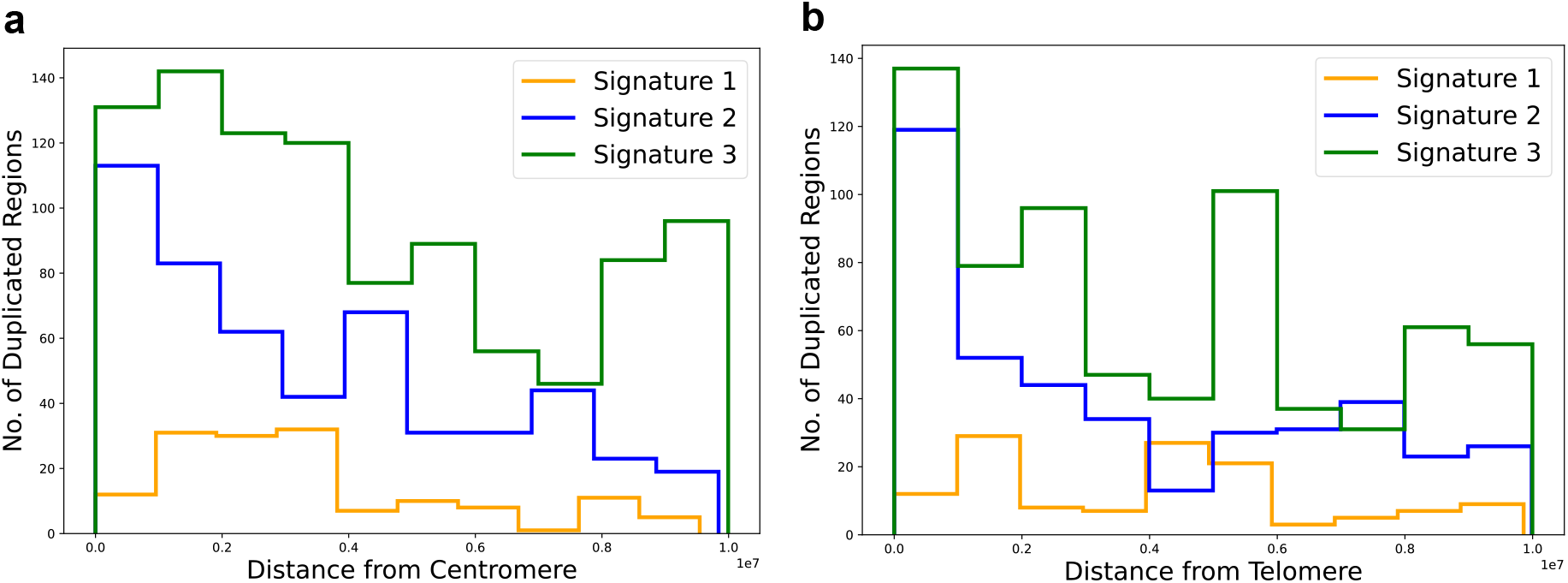
Distribution of duplicated regions assigned to three signatures relative to centromeres (**a**) and telomeres (**b**) of human chromosomes. Points represent non-overlapping sorted Mb windows. Leftmost point on both panels represents centromere or telomere proximal windows. One can see that s2 and s3 duplications are concentrated in subtelomeric and pericentromeric regions, while s1 is distributed uniformly.

By reducing original genomic feature space using an UMAP embedding we observed two clusters of duplicated regions McInnes et al. [2018]. These two clusters divided points according to a component size, where smaller cluster represented points from the giant component. Almost exclusively s2 points were located in the giant component cluster, while s1 and s3 represented rather clear subclusters of other points (Fig. 9a). Overall, the UMAP embedding seem to support our signatures assignment, because points belonging to same signatures tend to co-localize. Clustering of points according to a chromosome position was not as prominent. (Fig. 9b).

**Figure 9:**
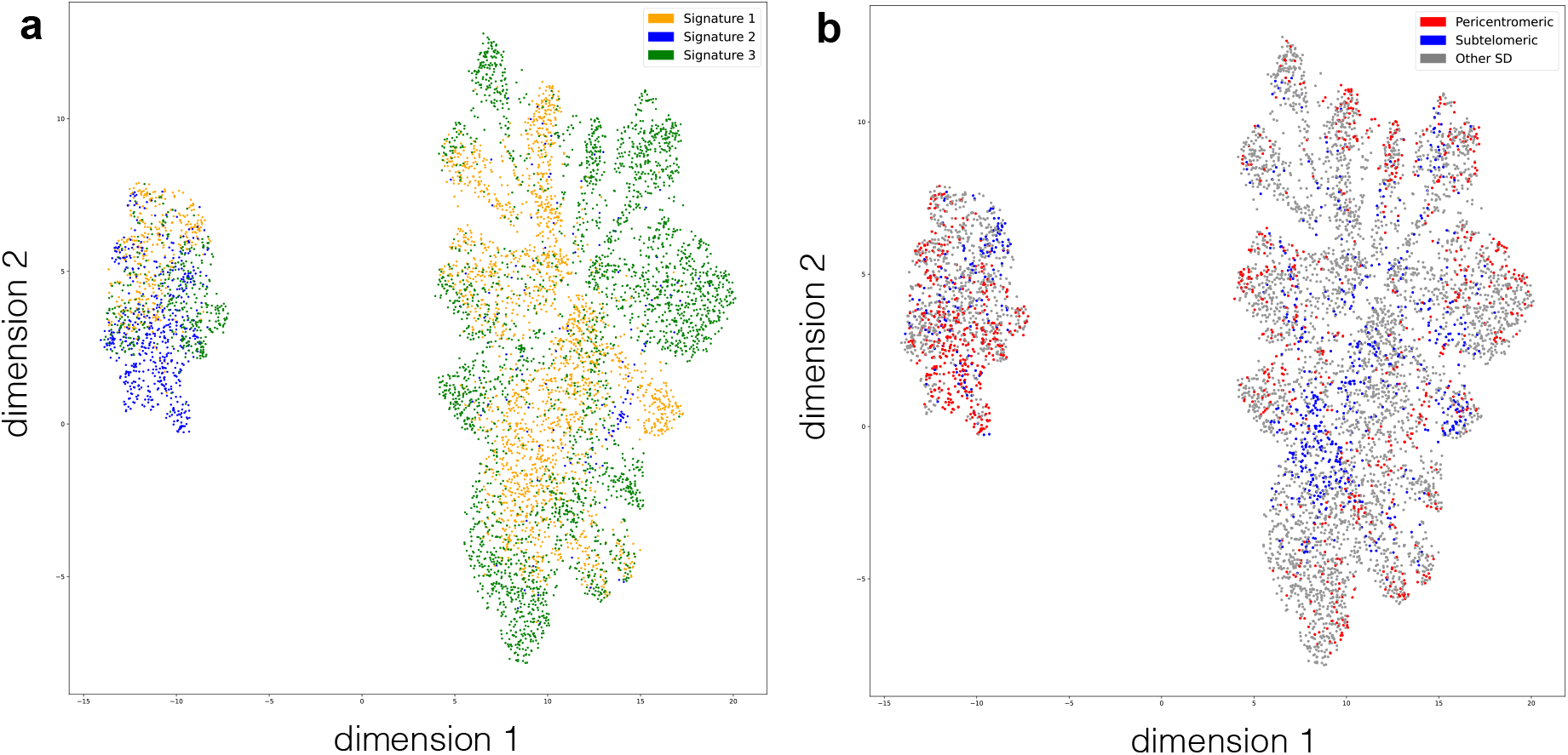
UMAP embedding of duplicated regions characteristics. Smaller cluster corresponds to points belonging to the giant component while larger one includes all others. Points are colored according to assigned signatures (**a**) or genomic positions (**b**). Points belonging to the same signature tend to co-localize. Specifically, s2 points are enriched in the giant component cluster.

## 3 Discussion

To study segmental duplications in the human genome we constructed the SD network. The nodes of it represent genomic regions involved in duplications and edges indicate the presence of an alignment between two corresponding regions. This network approach allows studying segmental duplications as a whole thus revealing some universal biological principles of SDs evolution from the SD network. Earlier we observed that duplication rates grow with the number of copies of a specific duplicated region, which we called preferential duplication rates Abdullaev et al. [2021]. This time we further examined the biological characteristics that could affect duplication rates.

We estimated the number of duplications that each node did during network growth. This was done in two steps. Firstly, we assigned edges weights inversely proportional to the number of shared neighbors. Secondly, we searched for the minimum spanning tree (MST) that goes through edges of minimal overall weight in order to discriminate real duplication events from secondary alignments. There is no *a priori* information on how many times each genomic region was duplicated in real life. Thus the algorithm was validated on PCM synthetic networks and some additional indirect evidence was used to estimate its quality. Manual analysis of duplicated regions (often a complicated task) can give more accurate reconstruction of corresponding duplication events, but it can not be used on a genome-wide scale.

We found that genomic characteristics are different in duplicated regions when comparing to the rest of the genome. It can be explained by highly non-random distribution of SDs in the human genome. For example, assembly gaps were enriched at duplicated regions breakpoints. This stems from the fact that complex duplication events are harder to assemble than non-redundant sequences, these ones are often not fully “embedded” in a genome assembly. The gene content was higher in duplicated regions which agrees with earlier report Zhang et al. [2005], moreover, it was positively correlated with the number of duplications of a region. This could be because actively duplicating regions are responsible for gene duplications and innovations. The number of overlapping CpG islands, to our knowledge, was not reported as associated feature, however, based on our analysis, it was both enriched in duplicated regions and positively correlated with duplication rates. We would like to note that even though CpG islands are correlated with both the G/C content and gene density, the partial correlation coefficients and the random forest importance values are not subjected to corresponding biases. We found that duplicated regions are located in late replicating parts, moreover, the DNA replication is slower in duplicated regions in comparison with the genome average and this affect is more prominent in actively duplicating loci. This evidence supports earlier report of Chen et al. [2015] and additionally suggests a new correlation with duplication rates. Recombination rates seem to be lower in duplicated regions, while G/C content is higher.

Another big group of features that we studied are high-copy repeats. We did not find strong associations between the composition of repeats and duplication rates. However, their composition is quite distinct in genomic windows located at different positions relative to breakpoints. Those repeat families depleted inside of duplicated regions and around the breakpoints, while their fraction is higher in the genome rest include: DNA transposons, *L1*, *L2*, *LTR* and *MIR* repeats. One can suggest that these repeats are rarely involved in duplication events. Another group includes repeats that are enriched at breakpoints thus, likely, making genome susceptible to duplications. These are satellite, microsatellite, *Alu* and low complexity repeats. It is unlikely that observed distributions relative to breakpoints are attributed to a non-uniform SDs distribution in the human genome (concentration of SDs in subtelomeric or subcentromeric regions, proximity to assembly gaps etc.). Firstly, in most cases frequencies of repeats change dramatically when moving short distances away from breakpoints which is not expected if only a biased placement of SDs play a role. Also, repeats distributions did not change substantially when we excluded duplicated regions proximal to assembly gaps.

Finally, we studied characteristic patterns (signatures) of a duplication process. We observed three signatures with various biological characteristics. Duplications belonging to signatures s2 and s3 are concentrated in pericentromeric and subtelomeric regions. s2 duplications are long and complex multi-copy loci which are overrepresented in the giant component of the SD network. These SDs are often flanked with assembly gaps, satellite repeats and non-autonomous retrotransposons. On the other hand, s3 duplications are much shorter, have much lower C/G content and are dominated with other groups of repeats at flanks: DNA transposons, *L1* and L2. The s1 group of duplications is the most interesting. These SDs are distributed in interstitial manner in regions of higher recombination rates with *Alu* and *MIR* repeats being enriched in their flanks. We suggest that these duplications likely originate from SINE-based non-allelic homologous recombination. These SDs are responsible for gene duplications and thus play an important evolutionary role. These observations support the hypothesis that the burst of *Alu* elements in human ancestral lineage gave a substrate for further interstitial duplications which often harbored genes and, as a result, lead to many human-specific gene innovations Bailey and Eichler [2006], Bailey et al. [2003]. We see a signature of this process predicted in unsupervised manner by our method.

## 4 Methods

### 4.1 Genomic data

The list of annotated SDs for the GRCh38 reference genome and all genomic features we considered in association tests were downloaded from the UCSC genome browser website (https://genome.ucsc.edu). For our analysis we disregard segmental duplications belonging to the sex chromosomes.

The genomic features we used include: replication timing, recombination rates, openness of chromatin, genome assembly gaps, CTCF sites, coordinates of a duplicated region, G/C nucleotides content, number of gene overlaps, repeat overlaps and CpG island overlaps (see Table 3). UCSC LiftOver tool was used to transfer coordinates from *hg19* to *hg38* where needed Kent et al. [2002]. The feature values were measured either inside of a duplicated region (between its breakpoints) or in flanking regions of length 50 bps padding the duplicated regions on both sides (“Position” column at Table 3). Some of the features are cell line specific, however, the cell lines where segmental duplications happened belong to the germline. In absence of a relevant data from the germline, we picked the source of the data as listed in the “Cell line” column. The feature types include: counts (repeats, assembly gaps etc.), mean values in corresponding genomic intervals (replication timing, recombination rate etc.) and fractions (fraction of G/C nucleotides, intrachromosomal edges from all neighbors of a node). Other than that, a span of replication timing, i.e. the difference between maximal and minimal timing values in a genomic interval, was measured and used as a proxy for the replication pausing (“Replication pausing” at Table 3). For those features where flanking regions are studied we did not distinguish between flanks and used the sum (or mean) of two values.

High-copy repeats distribution relative to duplicated regions breakpoints was studied by counting their numbers in sliding non-overlapping windows. These windows were first placed proximal to breakpoints of a duplicated region (coordinates: [0; 49] where 0 corresponds to a breakpoint position). Then moved 30 steps away ([0; 49], [50; 99],..., [1450; 1499]) and 5 steps inside of a duplicated region ([–250; –201], [–200; –151],..., [–50; –1]). These sliding windows moved symmetrically (as a pair) relative to breakpoints on both sides of a duplicated region and counted the number of repeat overlaps in each pair of windows. Moreover, to study the background distribution the sliding windows coordinates were randomly shuffled throughout the human genome 10 more times.

### 4.2 Network analysis

We generated a network of SDs where each node represents a duplicated region while edges connect nodes if an alignment between two corresponding duplicated regions exists. To each edge of the SD network we assigned a weight as described in the “Results” section. Those weights are reversely proportional to the fraction of neighbors that two nodes share. We run the Kruskal’s algorithm to reconstruct the minimum spanning tree on the weighted SD network Kruskal [1956].

To evaluate the accuracy of our approach we generated PCM synthetic networks. We simulated the PCM network growth and kept information on edges status (“primary” edges representing duplication events or “secondary” ones). All PCM simulations were done with the parameters inferred at (Abdullaev et al. [2021]): *δ* = 5.1 * 10^-4^, *f* = 0.47. We also considered other centrality measures assigned to nodes of the SD network (Table 2) and estimated an accuracy of each of them when applied to our task.

We applied the permutation tests when studied whether edges of interest (high sequence identity ones) are enriched in the MST. By doing 1000 sampling rounds from all edges of the SD network we generated a background distributions to compare with the real values.

### 4.3 Associations analysis

To predict genomic features associated with the number of duplications we used several machine learning algorithms. The applied algorithms include: linear regression, support vector regression (SVR), decision trees and random forest. All of them reached a similar quality of predictions with the maximal % of variance explained *R*^2^ = 30.5% observed for the random forest (estimated based on a 5-fold cross-validation). To estimate a statistical significance of RF feature importance values we used a special permutation test described by Altmann et al. [2010]. Multiple rounds of permutation of a response variable with consequent runs of random forest algorithm allow to approximate a null distribution of importance values for each feature when no interaction between a predictor and response variables exist. The resulting empirical p-values allow overcoming biases described in Altmann et al. [2010].

### 4.4 NMF: Non-negative Matrix Factorization

Non-Negative Matrix Factorization (NMF) is a matrix decomposition method. NMF decomposes an *n* * *f* matrix named *A*_(*n,f*)_ into a product of two matrices, namely *W*_(*n,k*)_ and *H*_(*k,f*)_. The matrix *W* is called the weight matrix, and the *H* matrix is called the signature matrix Paatero and Tapper [1994]. NMF computation is done with multiplicative iterative algorithm. The Kullback-Leibler divergence function was used as loss function. We normalized our data by scaling columns between 0 and 1 (the Min-Max scaler).

#### 4.4.1 Finding the number of signatures *k*

Various techniques are available to find the optimal number of signatures k. By experimenting with synthetic datasets whose k is *a priori* known, we picked the following methods: Fogel and Young’s volume-based method (FYV) and Stability of signatures predicted Fogel et al. [2007]. The variation of the FYV method implemented here computes the determinant of the signature matrix and the stability of signatures looks for the similarity of signatures computed for random initialization of signature and weight matrices. The FYV method predicts the best k value as 4. The stability of signatures predicts two as the best value. Hence we picked three as the ideal k value after observing the signatures predicted by NMF.

## 5 Abbreviations

SD: Segmental duplication
PCM: Preferential copying model
MST: Minimum spanning tree
CNV: Copy number variation
HR: Homologous recombination
NAHR: Non-allelic homologous recombination
NMF: Non-negative Matrix Factorization
FYV: Fogel and Young’s volume-based method

## Notes

### Competing Interest Statement

The authors have declared no competing interest.

