## Supplementary Materials for "Reconstruction of segmental duplication rates and associated genomic features by network analysis"

### Supplementary figures

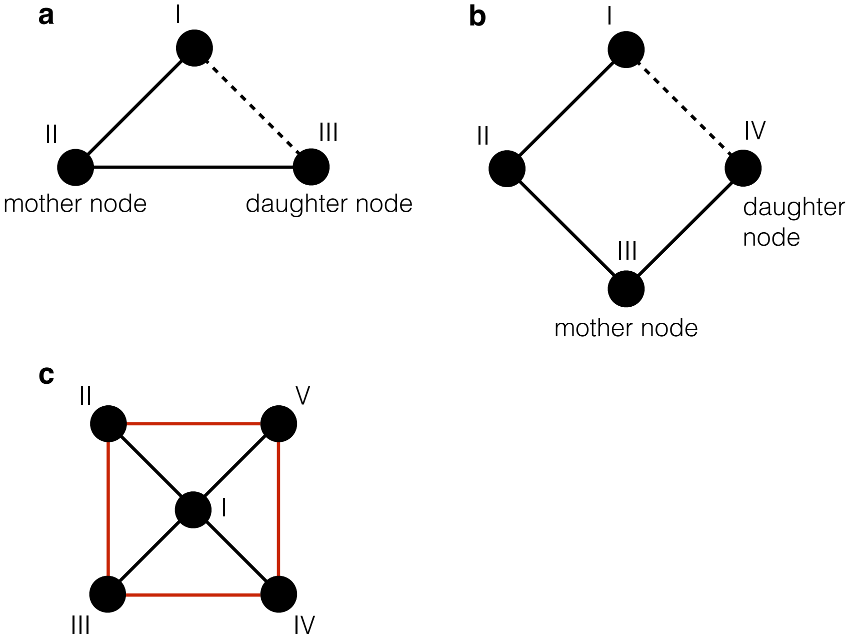


Sup. Fig. 1: Origin of cycles in the PCM. **a**. The scheme illustrates that cycles of size 3 can appear as a result of a duplication event when a daughter node inherits an edge from a mother node (the dotted line represents the secondary edge). **b**. On the other hand, cycles of size 4 or more can not appear in our copying model. The node $IV$ can not inherit the edge $(I,\ IV)$ from its mother node $III$, because the last one does not have the node $I$ among its neighbors. Thus we do not expect cycles of size 4 or larger in the network grown according to the PCM except for those cases when superposition of several cycles of size 3 forms cycles of larger size **c**. The cycle of size 4 (coloured in red) appears as a result of 4 cycles superposition: $(I,\ II,\ III)$, $(I,\ II,\ V)$, $(I,\ IV,\ V)$ and $(I,\ III,\ IV)$.

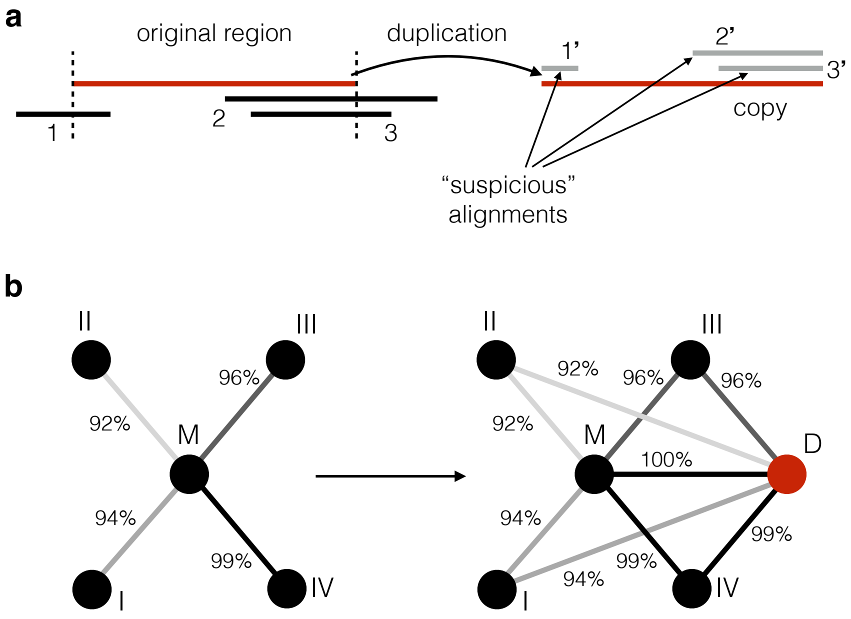


Sup. Fig. 2: The criteria we used to validated our predicted MST of duplications. The correct MST is depleted with suspicious alignments (nested ones with matching breakpoints). The scheme (**a**) illustrates this effect. The duplication event overlaps the alignments marked by 1, 2 and 3 and copies them incompletely (the loci to which 1, 2 and 3 align are not on the scheme). Thus the resulting copy aligns to the "mother" locus plus three additional loci (the alignments are marked as 1', 2' and 3' respectively) and since the original alignments were copied in abrupted manner, the alignments 1', 2' and 3' are suspicious according to our criterion. **b**. We suggest that at the moment of duplication two copies of a genomic sequence are almost identical (at least in a simple scenario of copy-paste process). Thus a "daughter" node inherits alignments (or secondary edges) with a sequence identity observed for corresponding alignments (edges) of a "mother" node which is illustrated on the scheme. Given a young age of high sequence identity edges, we expect them to be inherited more rarely than low identity ones. Thus we expect high sequence identity edges enrichment among primary edges.

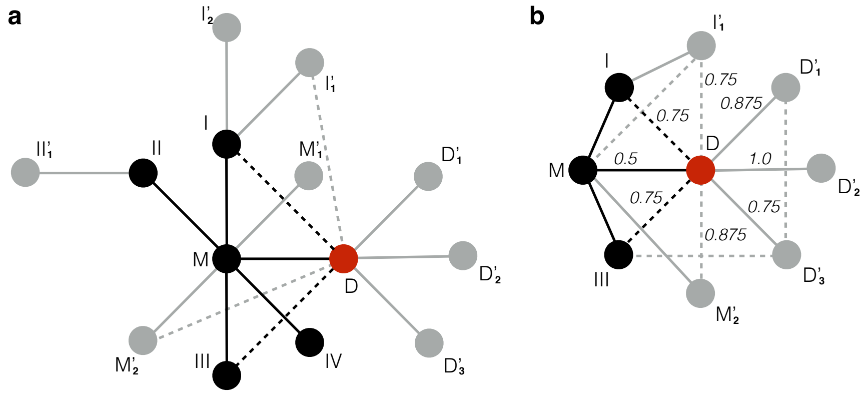


Sup. Fig. 3: **a**. All nodes in the nearest neighborhood of the node $D$ include: the mother node $M$, the neighbors of the mother node that were present before the birth of the node $D$ ($I,\ II,\ III,\ IV$), the daughters of the mother node that appeared after the birth of $D$ (denoted as $M_i^{'}$), the progeny of the nodes $I-IV$ denoted with apostrophes and the daughters of the node $D$ itself (denoted as $D_i^{'}$). The nodes and edges coloured in black were present at the moment of the birth of the node $D$, while gray ones appeared after. Dotted lines represent those secondary edges inherited by $D$ either at the moment of its birth (black ones) or when some other nodes were duplicated after the birth of $D$ (grey ones). For convenience we include on the first scheme only those secondary edges connected to $D$ while other ones are not shown. The scheme (**b**) includes the node $D$ and its neighborhood with all edges. On this scheme all secondary edges are illustrated. The edges weights are calculated as described in the text. As expected, the lowest weight is assigned to the edge connecting $D$ to its mother node $M$ because it shares the largest fraction of node $D$ neighbors.

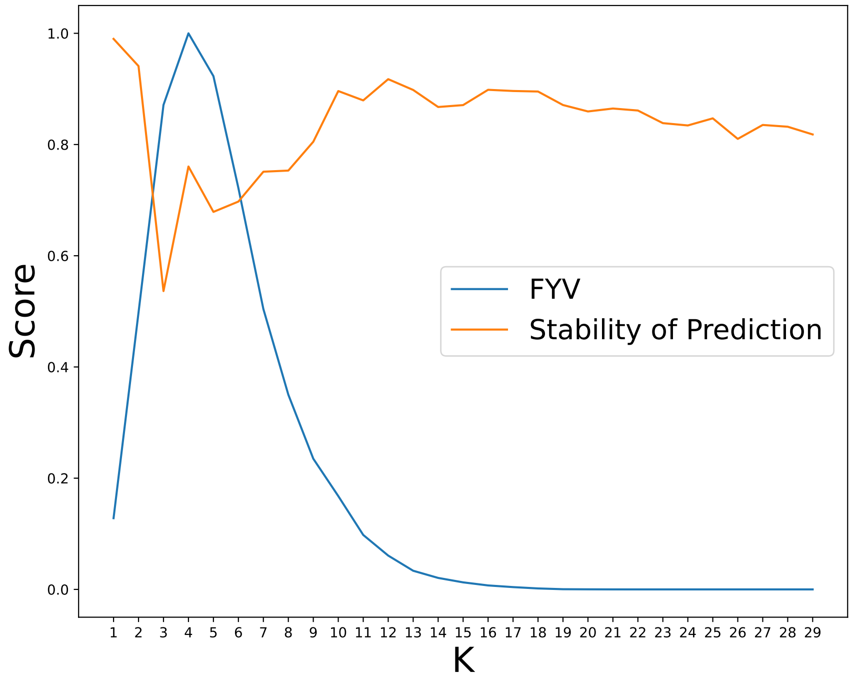


Sup. Fig. 4: Prediction of best signatures number $k$. The FYV method predicts the best $k$ value as 4. The stability of signatures predicts two as the best value. Eventually, we considered NMF with three signatures as an optimal solution.

### Additional validation of predicted MST

In a previous section we validated our method based on PCM simulations where we knew a correct order of duplication events. Here we wanted to use some indirect evidence to check if our MST predicted for the SD network gives a reasonable reconstruction of real duplication events. To do this we first applied the described algorithm to the SD network and checked if the resulting MST is enriched with real duplication events based on some features of edges in it.

If two or more alignments have their breakpoint coordinates matching, it is quite unlikely that this match happened as a random coincidence. This could happen because of some features of the sequence around the breakpoint (mechanic instability, NAHR hotspot etc.) or simply because this alignment is secondary (appears as a result of overlap of duplication events). One can find an illustration of how this matching breakpoints appear at the Sup. Fig. 2a. It means that an alignment nested inside of another longer alignment with one (or both) of the breakpoints matching between them is suspicious for being secondary. The longer alignment in a pair is, on the other hand, not suspicious (red alignment on the Sup. Fig. 2a). Not necessarily all secondary alignments are satisfying this criterion, but the alignment that satisfies it is likely secondary. In practice, we considered inexact matches where the distance between breakpoints is less than 5 bps. We collected all "suspicious" edges that correspond to alignments satisfying the described criterion and check if those edges are depleted in the MST predicted for the SD network. These "suspicious" edges with matching breakpoints were depleted among edges of our predicted MST in comparison with 1000 random samples of edges taken from the rest of the SD network (empirical p-value $< 0.001$).

Secondly, we expect that a correct MST covering real duplication events includes more edges of higher sequence identity than a random set of edges from the SD network. This might sound counter-intuitive, but edges with higher sequence identity are expected to be enriched in the correct MST of duplications in comparison with other secondary edges. In a simple scenario, we can assume that a level of sequence identity between two copies is the highest right after the moment of duplication and it decreases in a process of accumulation of neutral mutations with time. Secondly, when a duplicated region is duplicated again we expect that corresponding edge between "mother" and "daughter" node is of high sequence identity shortly after the duplication. On the other hand, edges inherited by a "daughter" node are of the same level of sequence identity as we observe for "mother" node (Sup. Fig. 2b). This can be illustrate with an example where a set of words represent duplicated regions while hamming distances are measures of sequence identity between them. If we make a copy of one of the words it will inherit all hamming distances to other words observed for the original word while the hamming distance between the original word and the new one will be equal to zero. Same behavior is expected for the sequence identity levels between duplicated regions. Thus at the moment when secondary edges of a "daughter" node appear, the level of their sequence identity is close to the one observed between the "mother" node and the corresponding neighbor. Primary edges, on the other hand, are of high sequence identity at the moment of their formation. This similarity drops in time in a process of mutation accumulation, however, we expect that highly identical alignments more likely correspond to real duplication events than those of lower sequence identity. We found that our predicted MST for the SD network is enriched with highly identical edges (alignments). There are more alignments of sequence identity higher than 0.99 in the predicted MST in comparison with random samples of edges from the rest of the SD network (empirical p-value = 0.001 based on 1000 permutation tests).

### Edge weight assignment

In this section we will discuss all types of node to node relationships and the fraction of shared neighbors that we expect. We can divide all potential neighbors of our node of interest $D$ into five categories based on their origin relative to it (Sup. Fig. 4). The first two categories include those nodes that were already present at the moment of $D$ birth and could potentially become its neighbors at that moment. These include one mother node and its neighbors at the moment of $D$ node birth (nodes $M$ and $I$, $II$, $III$, $IV$ at Sup. Fig. 4a respectively). The other three categories include those nodes that appeared after the moment of $D$ birth: the progeny of the mother node that appeared after the birth of our node (nodes $M_1^{'}$, $M_2^{'}$), the progeny of the neighbors of our node that appeared after the birth of our node (nodes $I_1^{'}$, $I_2^{'}$, $II_1^{'}$) and all the progeny of our node $D$ itself (nodes $D_1^{'}$, $D_2^{'}$ and $D_3^{'}$). Also, for convenience, several functions were introduced to define sets of nodes: $\mathcal{N}()$ is a set of neighbors of a specific node, $\mathcal{N}_{birth}()$ has the same meaning as $\mathcal{N}()$, but the set includes only those neighbors that were present at the moment of $D$ node birth, $\mathcal{P}()$ is a progeny of a specific node that appeared after the $D$ node birth. Let's give several examples of those functions use:

$$\begin{aligned}
\mathcal{N}(M) &= \{I, II, III, IV, D, M_1^{'}, M_2^{'}\}, \\
\mathcal{N}_{birth}(M) &= \{I, II, III, IV\}, \\
\mathcal{P}(M) &= \{M_1^{'}, M_2^{'}\}, \\
\mathcal{N}(I) &= \{M, D, I_1^{'}, I_2^{'}\}, \\
\mathcal{N}_{birth}(I) &= \{M\}, \\
\mathcal{P}(I) &= \{I_1^{'}, I_2^{'}\}.
\end{aligned}$$

As one will see in further calculations, on average, a mother node of any chosen node shares the highest fraction of its daughter node neighbors. We used this observation in our approach of assigning weights to edges of the SD network. For each edge $e_i$ between our node $D$ and $i^{th}$ neighbor $n_i$ we assign the weight $w_i = 1 - \frac{\|\mathcal{N}(D) \cap \mathcal{N}(n_i)\|}{k_D}$, where $k_D$ is a node degree of $D$ while $\|\cdot\|$ brackets denote the number of elements in a set (Sup. Fig. 4b). Values of $w_i$ lie in the interval $[0 < w_i \leq 1]$ where $w_i = 1$ when no neighbors of $D$ are shared with $n_i$, while $w_i > 0$ because at least one neighbor of $D$ ($n_i$ itself) is never among neighbors of $n_i$.

Now we can estimate the expected number of shared neighbors between $n_i$ and $D$, i. e. $\|\mathcal{N}(D) \cap \mathcal{N}(n_i)\|$ for neighbors belonging to all five described categories. For simplicity, in the last three categories which include neighbors inherited after the birth of $D$ only "nearest" progeny (daughter nodes) will be included (because "further" progeny shares even less neighbors with $D$). Let's start by defining the following sums:

$$\mathbf{I} = \sum_{i \in \mathcal{N}_{birth}(M)} 1(i)$$

The indicator function 1() here and in all formulas below is defined as: $1(i) = 1$ if $i$ is among neighbors of $D$ and $1(i) = 0$ otherwise.

$$\mathbf{P} = \sum_{i \in \mathcal{P}(I)} 1(i) + \sum_{j \in \mathcal{P}(II)} 1(j) + ... + \sum_{l \in \mathcal{P}(IV)} 1(l)$$

where $\mathbf{P}$ represents the number of nodes that are connected to $D$ among the after birth progeny of $\mathcal{N}_{birth}(M)$ nodes (i.e. among $I_1^{'}$, $I_2^{'}$, $II_1^{'}$ nodes in our case).

$$\mathbf{M} \;=\; \sum_{i \in \mathcal{P}(M)} 1(i)$$

where $\mathbf{M}$ represents the number of nodes that are connected to $D$ among $M_i^{'}$ (the progeny of $M$ that appeared after the birth of $D$).

$$\mathbf{D} \;=\; \sum_{i \in \mathcal{P}(D)} 1(i) \;=\; \|D^{'}\|$$

where $\mathbf{D}$ represents the number of nodes that are connected to $D$ among $D_i^{'}$ (or equivalently among $\mathcal{P}(D)$ ). This value just equals to the number of $D^{'}$ nodes.

Then an expected number of shared neighbors $\|\mathcal{N}(D) \cap \mathcal{N}(n_i)\|$ for neighbors $n_i$ belonging to all five categories of nodes described above can be calculated this way:

$$\begin{aligned}
\|\mathcal{N}(D) \cap \mathcal{N}(M)\| &= \mathbf{I} + f\mathbf{P} + \mathbf{M} + f\mathbf{D} \\
\|\mathcal{N}(D) \cap \mathcal{N}(M_i^{'})\| &= 1 + f\mathbf{I} + f^2\mathbf{P} + f(\mathbf{M} - 1) + \hat{f}\mathbf{D} \\
\|\mathcal{N}(D) \cap \mathcal{N}(l)\| &= 1 + \dot{f}(\mathbf{I} - 1) + \dot{f}f(\mathbf{P} - \sum_j 1(l_j^{'})) + \sum_j 1(l_j^{'}) + f\mathbf{M} + f\mathbf{D} \\
\|\mathcal{N}(D) \cap \mathcal{N}(l_i^{'})\| &= 1 + \dot{f}f(\mathbf{I} - 1) + \dot{f}f^2(\mathbf{P} - \sum_j 1(l_j^{'})) + f\sum_j 1(l_j^{'}) + f^2\mathbf{M} + \hat{f}\mathbf{D} \\
\|\mathcal{N}(D) \cap \mathcal{N}(D_i^{'})\| &= f\mathbf{I} + \hat{f}\mathbf{P} + \hat{f}\mathbf{M} + f\mathbf{D}
\end{aligned}$$

where $l \in \mathcal{N}_{birth}(M)$, $\dot{f}$ – is a probability of an edge being present between two nodes from the $\mathcal{N}_{birth}(M)$ set. It is proportional to the local clustering coefficient. The value $\hat{f} \in [f^2, f]$ and depends on the order of duplication events (for example, whether the node $I_1^{'}$ appeared before or after the $D_1^{'}$). One can check the correctness of these equations by going over all possible pairwise relationships between node types (which is $5 * 5 = 25$ pairs) and calculate the probabilities of sharing neighbors from corresponding node categories. These probabilities would represent the coefficients associated with each term of the above sums. For example, if we consider only $\|\mathcal{N}(D) \cap \mathcal{N}(M)\|$ equation, the mother node $M$ shares all neighbors of $D$ from the set $\mathcal{N}_{birth}(M)$ because all these nodes are connected to $M$ by definition; shares all neighbors of $D$ from $\mathcal{P}(M)$ set because $M$ is their mother node and thus connected to all of them;

shares fraction $f$ of $\mathcal{P}(D)$ nodes because these nodes inherit the edge to $M$ each with probability $f$ and, finally, shares the fraction $f$ of neighbors $D$ belonging to the $\{\mathcal{P}(I),\ \mathcal{P}(II),\ \mathcal{P}(III),\ \mathcal{P}(IV)\}$ set, because each node in it inherits the edge to $M$ from its mother node with probability $f$.

Now one can see that a mother node is expected to share the highest number of neighbors with a daughter node in comparison with other nodes. However, the bias could appear if one of the neighbors $n_i$ is actively duplicated in unbalanced manner after the birth of $D$. This, in our equations, means a high value of $\sum_{i \in \mathcal{P}(l)} 1(i)$ sum which inflates the $\|\mathcal{N}(D)\ \cap\ \mathcal{N}(l)\|$ value where $l \in \mathcal{N}_{birth}(M)$.

So the overall logic is the following: we know (from the PCM) that a mother node, on average, shares the highest number of neighbors of a daughter node in comparison with other nodes in a neighborhood of a daughter node. Thus after assigning weights in the described manner we expect that edges connecting mother and daughter nodes would, on average, have lower weights, thus minimum spanning tree covering all nodes in the graph and going through edges of minimal overall weight would be enriched with primary alignment edges (as opposed to secondary alignment edges) that represent real duplication events. Finally, let's note that an edge weight depends on a node we pick: for an edge $e$ between vertices $a$ and $b$ the weight $w_e$ can be calculated as $w_e = 1\ -\ \frac{\|\mathcal{N}(a)\ \cap\ \mathcal{N}(b)\|}{k_a}$ or $w_e = 1\ -\ \frac{\|\mathcal{N}(a)\ \cap\ \mathcal{N}(b)\|}{k_b}$ and those values are ordinarily not the same. In practice we assigned the least of two values as a weight of specific edge which equals $w_e = 1\ -\ \frac{\|\mathcal{N}(a)\ \cap\ \mathcal{N}(b)\|}{min(k_a,\ k_b)}$.
